# The Small Tumor Antigen of Merkel Cell Polyomavirus Accomplishes Cellular Transformation by Uniquely Localizing to the Nucleus Despite the Absence of a Known Nuclear Localization Signal

**DOI:** 10.1101/2023.11.28.569067

**Authors:** Kaira R. Thevenin, Isabella S. Tieche, Cody E. Di Benedetto, Matt Schrager, Kristine N. Dye

**Author notes:** Corresponding Author: Kristine N. Dye; 421 N Woodland Blvd, DeLand, FL, 32723.

## Abstract

**Background:** Merkel Cell Carcinoma (MCC) is an aggressive skin cancer that is three times deadlier than melanoma. In 2008, it was found that 80% of MCC cases are caused by the genomic integration of a novel polyomavirus, Merkel Cell Polyomavirus (MCPyV), and the expression of its small and truncated large tumor antigens (ST and LT-t, respectively). MCPyV belongs to a family of human polyomaviruses; however, it is the only one with a clear association to cancer.

**Methods:** To investigate the role and mechanisms of various polyomavirus tumor antigens in cellular transformation, Rat-2, 293A, and human foreskin fibroblasts were transduced with pLENTI MCPyV LT-t, MCPyV ST, TSPyV ST, HPyV7 ST, or empty pLENTI and assessed through multiple transformation assays, and subcellular fractionations. One-way ANOVA tests were used to assess statistical significance.

**Results:** Soft agar, proliferation, doubling time, glucose uptake, and serum dependence assays confirmed ST to be the dominant transforming protein of MCPyV. Furthermore, it was found that MCPyV ST is uniquely transforming, as the ST antigens of other non-oncogenic human polyomaviruses such as Trichodysplasia Spinulosa Polyomavirus (TSPyV) and Human Polyomavirus 7 (HPyV7) were not transforming when similarly assessed. Identification of structural dissimilarities between transforming and non-transforming tumor antigens revealed that the uniquely transforming domain(s) of MCPyV ST are likely located within the structurally dissimilar loops of the MCPyV ST unique region. Of all known MCPyV ST cellular interactors, 62% are exclusively or transiently nuclear, suggesting that MCPyV ST localizes to the nucleus despite the absence of a canonical nuclear localization signal. Indeed, subcellular fractionations confirmed that MCPyV ST could achieve nuclear localization through a currently unknown, regulated mechanism independent of its small size, as HPyV7 and TSPyV ST proteins were incapable of nuclear translocation. Although nuclear localization was found to be important for several transforming properties of MCPyV ST, some properties were also performed by a cytoplasmic sequestered MCPyV ST, suggesting that MCPyV ST may perform different transforming functions in individual subcellular compartments.

**Conclusions:** Together, these data further elucidate the unique differences between MCPyV ST and other polyomavirus ST proteins necessary to understand MCPyV as the only known human oncogenic polyomavirus.

## BACKGROUND

Merkel Cell Carcinoma (MCC) was first described in the 1970s as a rare, aggressive skin cancer, likely a consequence of ultraviolet radiation-induced mutations in cell cycle-regulation genes (1–3). However, in 2008, a novel human polyomavirus, subsequently termed Merkel Cell Polyomavirus (MCPyV), was found to be integrated within, and the etiologic agent of, ∼80% of MCC tumors (4). Further investigation identified MCPyV as a ubiquitous, asymptomatic human virus that, in rare cases of immunosuppressed individuals, could integrate into the human genome and lead to the development of cancer (5–9).

The small, 5.3kb dsDNA genome of MCPyV is divided into an early region and a late region (ER and LR, respectively) by a bidirectional promoter (4,10). Due to its constrained genomic size, the ER utilizes alternative splicing and overprinting to generate 4 viral proteins, including the large tumor antigen (LT), small tumor antigen (ST), alternate large T open reading frame (ALTO), and the 57kT protein, while the LR encodes the VP1 and VP2 structural proteins (10–12). Although all of these viral proteins are expressed for successful viral replication in the normal MCPyV viral lifecycle, in MCC, only a truncated LT (LT-t) and ST are expressed and necessary for the viability of MCC tumors, suggesting a role for these proteins in tumorigenesis and maintenance (4,13,14). Several researchers have identified transforming capabilities of both ST and LT-t, including, but not limited to, perturbation of MYCL and Rb, respectively (10,15–19).

MCPyV belongs to a family of 14 known human polyomaviruses but is currently the only one with a clear association to cancer (20,21). However, polyomaviruses of other species, such as Simian Virus 40 (SV40), which infects simian species, has been found to be oncogenic in rodents (22–25). As one of the most intensively studied oncogenic viruses, the study of SV40 oncogenesis has led to the development of several transformation assays, which have subsequently been used to direct the study of MCPyV transformation and tumorigenesis.

Transformation is defined as the acquisition of expanded proliferation and/or survival potential of a cell (26). Many *in vitro* transformation assays exist, including soft agar assays, focus formation, doubling time, proliferation rate, serum dependence, and metabolic assays, each of which assesses different aspects and pathways involved in cellular transformation that may collectively contribute to tumorigenesis *in vivo.* In order to accomplish cellular transformation, viral proteins bind and perturb cellular pathways that allow for expanded proliferation and survival potential (27,28). Cellular proteins and pathways operate in specific subcellular compartments, including the cytoplasm, nucleus, and membrane (29,30). As localization influences the regulation and activity of specific proteins and pathways, it is tightly regulated (30,31). Therefore, protein localization to the nucleus is tightly regulated and restricted to proteins containing a nuclear localization signal (NLS), as this is where the DNA is contained, and gene expression is initiated (32).

Herein, we employed an alternative approach to elucidate the mechanism of transformation of MCPyV T antigens. As opposed to previous approaches of comparing similarities between SV40 and MCPyV T antigens, several transformation assays on various oncogenic and non-oncogenic human polyomavirus tumor antigens were performed to identify the unique domains and mechanisms of MCPyV T-antigen mediated oncogenesis. Together, we identified MCPyV ST to be predominantly and uniquely transforming when compared to MCPyV LT-t and the ST of other, skin-tropic human polyomaviruses. Furthermore, MCPyV ST was found to uniquely localize to the nucleus, consistent with its known interaction with cellular nuclear proteins, despite the absence of a canonical NLS. This is in contrast to non-oncogenic human polyomavirus ST proteins, which localize exclusively to the cytoplasm. The nuclear localization of MCPyV ST was found to be responsible for many, but not all, MCPyV ST transforming properties, suggesting MCPyV ST binds to and perturbs many pathways in both the cytoplasm and nucleus to accomplish cellular transformation. Together, these data further increase our understanding of MCPyV ST-mediated cellular transformation and may prove influential in further discoveries of MCPyV ST-mediated oncogenesis and the consequent development of MCPyV-targeted MCC therapies.

## METHODS

### Plasmids and Mutagenesis

Codon-optimized constructs with a Kozak sequence were subcloned into pLENTI-puro (Addgene #39481) to create MCPyV ST ko/co, TSPyV ST ko/co, and HPyV7 ST ko/co, which were transduced into Rat-2 cells. pLENTI-puro empty and pLENTI-puro MCPyV LT-t were also transduced into Rat-2 cells. The Q5 NEB Mutagenesis Kit (New England Biolabs) was used to add an NES (LQKKLEELEL) or NLS (SPKKKRKVE) followed by a (GGGGS)_2_ flexible linker to the 5’ end (N-terminus) of pLENTI-puro MCPyV ST ko/co to create pLENTI-puro NES-MCPyV ST ko/co and pLENTI-puro NLS-MCPyV ST ko/co. pMTBS (donated by Dr. Christopher Buck, NIH) was used to transfect 293A cells with MCPyV ST. TSPyV ST and HPyV 7 ST were subcloned into pCS2 to transfect 293A cells. Empty pLENTI and pCS2 were used as negative controls.

### Tissue Culture, Transductions, and Transfections

Adenovirus-transformed human embryonic kidney cells (HEK293A), 293TNs, and Rat-2 cells were cultured in 1x DMEM (ThermoFisher Scientific) supplemented with 10% fetal bovine serum (FBS) (Neuromics), GlutaMax (ThermoFisher Scientific), Non-Essential Amino Acids (NEAA) (ThermoFisher Scientific), and 100 units/mL penicillin-streptomycin (ThermoFisher Scientific). MCC virus-positive MKL-1 cells were cultured in 1x RPMI (ThermoFisher Scientific) supplemented similarly as stated above. All cell lines were incubated at 37°C in 5% CO_2_.

293TN cells were transfected with the lentiviral envelope and packaging plasmids psPAX2 (Addgene #12260) and pMD2.G (Addgene #12259) and lentiviral transfer plasmid pLENTI-puro using TransIT-293 Transfection reagent (Mirus). Viral harvests were conducted at 24 and 48 hours post-transfection. Rat-2 cells were transduced with the pseudotyped lentiviruses supplemented with 4μg/mL polybrene (SantaCruz Biotechnology) and selected with 4μg/mL puromycin (ThermoFisher Scientific) 48 hours post-transduction.

HEK293 cells were transfected with pMTBS and pCS2-ST at ∼80% confluence using TransIT-293 transfection reagent (Mirus). Cells were harvested for downstream applications 48 hours post-transfection.

### Immunoblotting and Antibodies

Cell lysates were prepared in RIPA lysis buffer (ThermoFisher Scientific) supplemented with a protease inhibitor cocktail (ThermoFisher Scientific) on ice, followed by sonication at 20% amplitude and centrifugation at 16,000 x g for 15 minutes. A BCA assay (ThermoFisher Scientific) was performed to determine protein concentrations for normalization in 2x SDS sample buffer (ThermoFisher Scientific) at 1μg/μL followed by boiling at 95°C. 30μg of protein was loaded in an 8-16% SDS-PAGE gel (ThermoFisher Scientific) and transferred to an Immobilon-P PVDF membrane, followed by a 1 hour block in 4% milk and overnight primary antibody incubation with Ab5 (donated by James Decaprio, Harvard University) diluted at 1:1000 for MCPyV ST, HPyV7 ST, and TSPyV ST, and MCPyV LT, HSP90 (Cell Signaling Technologies, 4874S) (1:5000), HDAC2 (Cell Signaling Technologies, 2540S) (1:5000), Na/K-ATPase (Cell Signaling Technologies, 3010S) (1:2500), or actin (Cell Signaling Technologies, 5125S) (1:10,000). After TBST washes, membranes were incubated in 4% milk supplemented with either anti-mouse IgG HRP-conjugated secondary antibody (Cell Signaling Technologies, 7076P2) (1:10,000) or anti-rabbit IgG HRP-conjugated secondary antibody (Cell Signaling Technologies, 7074P2) (1:10,000) at room temperature for 1 hour. Following secondary washing, proteins were detected using Femto (ThermoFisher Scientific) and a ChemiDoc Imaging System (Bio-Rad).

### Transformation Assays

Soft agar assays were performed in triplicate according to the JOVE online protocol at 50,000 cells/well (33). The top layer of 1x DMEM was supplemented with puromycin. The cells were monitored for three weeks, followed by imaging with a Nikon Ti2 Eclipse microscope (Nikon).

Metabolism experiments were performed in triplicate using a Glucose Uptake-Glo Assay Kit (Promega) according to manufacturer instructions at 10,000 cells/96-well.

Serum dependence experiments were performed by plating 50,000 cells/6-well in triplicate in complete DMEM supplemented with either 10%, 5%, or 1% FBS (Neuromics). Cells were quantified at days 2, 5, and 10. Puromycin-supplemented media was changed every 3 days or as needed in later time points.

Proliferation rate experiments were conducted by plating 50,000 cells/6-well in triplicate. Cells were quantified at days 2, 4, 8, 14, 20, and 28. Doubling time was evaluated between days 0 and 28 to be representative of the entire experiment. Puromycin-supplemented media was changed every 3 days or as needed in later time points.

### Subcellular Fractionation

Subcellular fractionations were conducted using a Subcellular Fractionation Kit (ThermoFisher Scientific) following a manufacturer-modified protocol. In short, cells were plated at 1×10^6^ in 4 10cm plates 24 hours prior to harvest through trypsinization (ThermoFisher Scientific) for adherent cells. Provided fractionation extraction buffers were supplemented with a 1x protease inhibitor cocktail and used to fractionate the cell sample as instructed with the addition of extra wash steps to lessen fraction contamination. Following protein extraction, lysates were quantified and run in a western blot as described previously. Controls included Heat Shock Protein 90 (HSP90) for the cytoplasmic fraction, Sodium Potassium ATPase (NA/K ATPase) for the membrane fraction, and Histone Deacetylase 2 (HDAC2) for the nuclear soluble fraction.

### Statistical Methods

Statistical analyses were conducted using the Jamovi statistical computer software, Version 2.3. All experiments were repeated in triplicate. One-way ANOVA analyses were used, and a *p-value* <0.05 was considered to indicate a statistically significant difference.

## RESULTS

### The ST unique region is responsible for the dominant transforming properties of MCPyV ST

In order to investigate the mechanism(s) of MCPyV cellular transformation, we first sought to assess various transforming capabilities of the MCPyV tumor antigens (TAgs) expressed in virus positive MCC (VP-MCC): ST and LT-t. As done previously by others, Rat-2 cells were transduced with pseudotyped lentiviruses containing plasmids encoding MCPyV ST, LT-t, or empty pLENTI control (34). After confirmation of protein expression (Fig. 1A), soft agar assays were performed to assess the ability of the MCPyV TAgs to induce anchorage-independent cellular proliferation, a common property of cancerous cells that promotes tumor formation and metastasis (35). Consistent with the work of others, MCPyV ST was capable of inducing anchorage-independent growth, whereas MCPyV LT-t was incapable of colony formation (Fig. 1B and C) (34,36).

**Figure 1.**
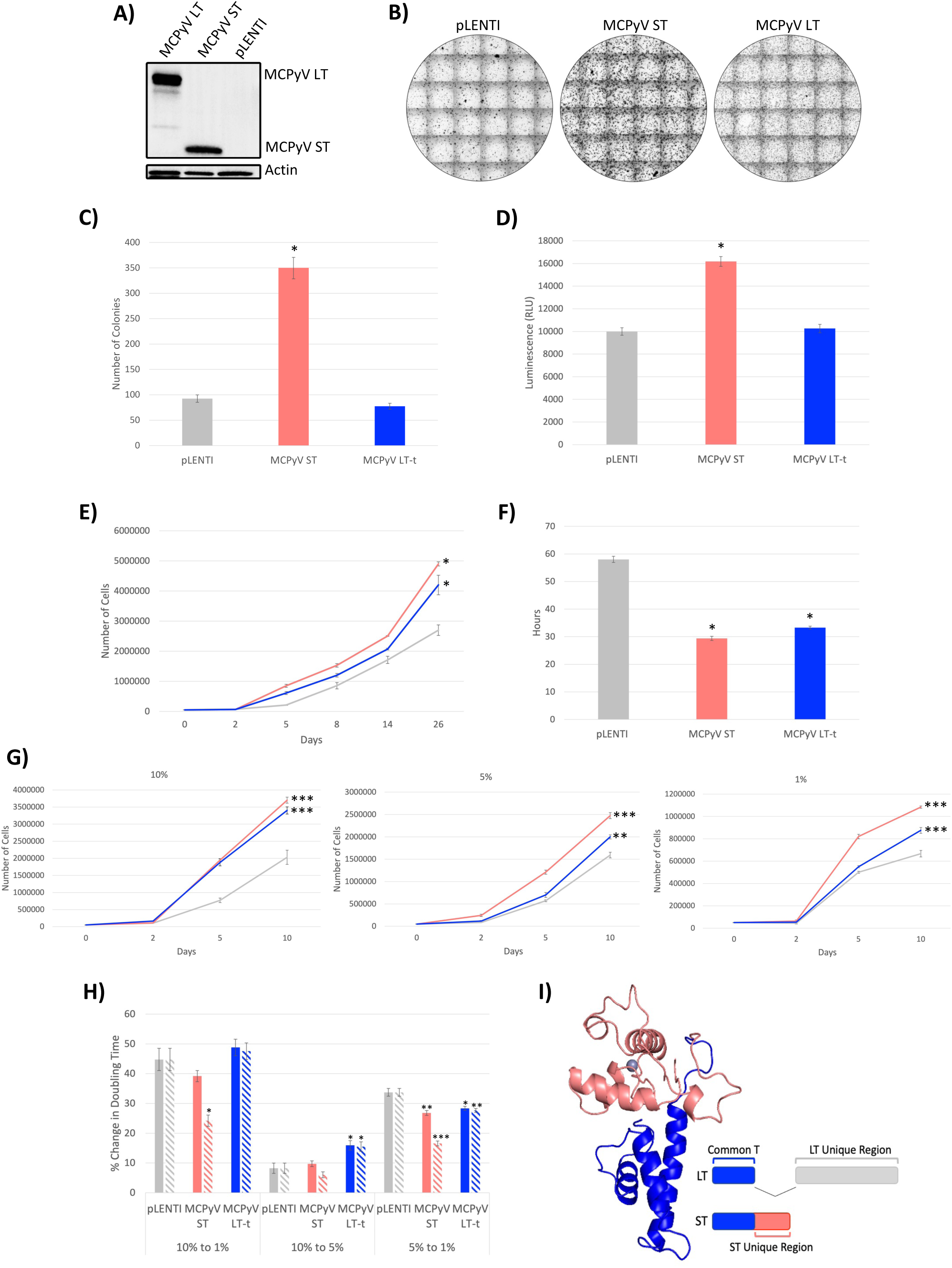
The ST unique region is responsible for the dominant transforming properties of MCPyV ST. Rat-2 cells were transduced with MCPyV LT-t, MCPyV ST, or pLENTI control and protein expression was confirmed (A). Various transformation assays were performed including soft agar assays (B and C), glucose uptake (D), proliferation rate (E), doubling time (F), and serum dependence (G) (grey – pLENTI, coral – MCPyV ST, blue – MCPyV LT-t). The percent change in serum dependence doubling time (solid bars) was normalized (dashed bars) by glucose uptake compared to pLENTI control (H). MCPyV ST protein structure was determined by PyMol and colored to show the common-T (blue) and ST unique (pink) regions (I). Each point represents mean ± the standard error of independent triplicates. One-way ANOVA * p < 0.05; ** < 0.01; *** p < 0.001, significantly different from pLENTI.

Because cellular transformation can be defined by properties other than anchorage-independent growth, additional transformation assays were performed between the MCPyV TAgs to further define the specific transforming properties of both MCPyV ST and LT-t. It has been frequently described that cancerous cells have an increased metabolic rate and, therefore, glucose uptake than healthy cells to support their robust cellular division (37,38). Consistently, MCPyV ST expression significantly increased the glucose uptake of Rat-2 cells, whereas MCPyV LT-t did not (Fig. 1D). Similarly, MCPyV ST was also capable of significantly increasing the proliferation rate and decreasing the doubling time of Rat-2 cells (Fig. 1E and F); however, these assays demonstrated that MCPyV LT-t was also capable of significantly increasing the cellular proliferation rate and decreasing the doubling time compared to control cells, albeit to a lesser degree than MCPyV ST. These data support the transforming properties of both MCPyV ST and LT-t, consistent with the necessity of both ST and LT-t to be expressed for the viability of VP-MCC cells.

Finally, the serum dependence of Rat-2 cells was compared between control, ST, and LT-t expressing cells, as cancerous cells are frequently described as capable of maintaining cellular proliferation in the absence of the growth factors and nutrients found in serum (39). Consistent with proliferation rate experiments containing the normal 10% serum supplementation (Fig. 1E), both ST and LT-t were capable of significantly increasing proliferation in 10% serum; however, MCPyV LT-t exhibited less proliferation than MCPyV ST with reduced serum concentrations of either 5% or 1% (Fig. 1G). To illustrate the effect of reducing serum concentrations on the proliferation of the control, ST, and LT-t expressing cells, the percent change in doubling time was calculated going from 10% to 1%, 10% to 5%, and 5% to 1% serum. Although the percent change in doubling time between these different serum concentrations was not significantly different between control, ST, and LT-t expressing cells when going from 10% to 1% serum, normalization by glucose uptake (Fig. 1D) found MCPyV ST to significantly decrease the percent change in doubling time upon serum reductions to 1% (Fig. 1H). Together, these data found serum concentrations to affect the proliferation of ST-expressing cells the least, despite these cells having the highest metabolic rate as inferred by glucose uptake.

Upon confirmation of ST as the dominant transforming protein of MCPyV, we sought to elucidate the domain(s) responsible for this phenotype. Both MCPyV ST and LT-t share the same start codon and are created through the alternative splicing of the same mRNA (11). Therefore, both ST and LT-t share the same N-terminal amino acid sequence, referred to as “common-T”, whereas they differ in their C-terminal amino acid sequence, which is referred to as the “ST- and LT – unique regions” (Fig. 1I). The finding of ST to be the dominant transforming protein of MCPyV led to the hypothesis that the domain(s) responsible for these unique transforming properties are likely located in the ST-unique region, as location within the common-T region would suggest MCPyV LT-t to have similar transforming properties. Together, these data confirm ST as the dominant transforming protein of MCPyV by assessment of various transformation properties such as anchorage-independent growth, metabolism, proliferation rate, doubling time, and serum dependence, and that the domain(s) responsible for this phenotype are likely found in the ST unique region.

### The structurally dissimilar loops in the ST unique region of MCPyV ST may be responsible for the uniquely transforming capabilities of MCPyV ST when compared to other skin-tropic human polyomaviruses

Upon confirmation of ST being the dominant transforming protein of MCPyV, likely a result of activities in the ST unique region, we sought to further elucidate the region(s) of MCPyV ST responsible for various transformation properties. As noted, MCPyV ST belongs to a family of 14 known human polyomaviruses, but is the only polyomavirus found to be directly responsible for the development of cancer (20,21). Therefore, we sought to determine whether the ST antigens of other, skin tropic, human polyomaviruses, such as Human Polyomavirus 7 (HPyV7) and Trichodysplasia Spinulosa Polyomavirus (TSPyV), were also capable of any of the mechanisms of cellular transformation similar to MCPyV ST. Rat-2 cells were transduced to express MCPyV, TSPyV, or HPyV7 ST (Fig. 2A). Although it appeared that both HPyV7 and TSPyV ST proteins were expressed in lower quantities than MCPyV ST, it is likely that the antibody used to detect these proteins has higher affinity to MCPyV ST, as it was developed through its recognition of the MCPyV ST antigen; however, each T antigen was similarly codon optimized and under the control of the same CMV promoter.

**Figure 2.**
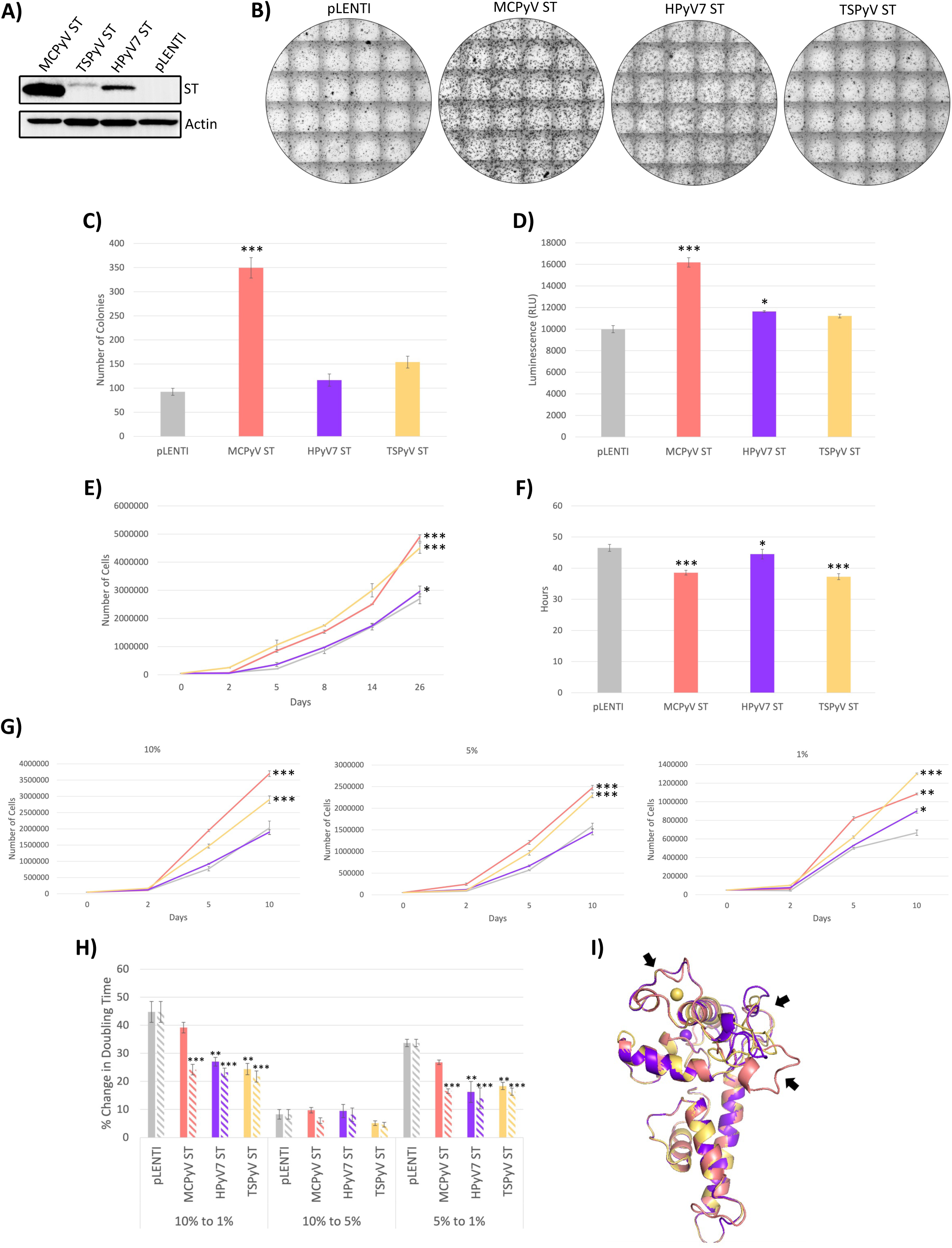
The structurally dissimilar loops in the ST unique region of MCPyV ST may be responsible for the uniquely transforming capabilities of MCPyV ST when compared to other skin-tropic human polyomaviruses. Rat-2 cells were transduced with MCPyV ST, HPyV7 ST, TSPyV ST, or pLENTI control, and protein expression was confirmed (A). Various transformation assays were performed, including soft agar assays (B and C), glucose uptake (D), proliferation rate (E), doubling time (F), and serum dependence (grey – LENTI, coral – MCPyV ST, purple – HPyV7 ST, yellow – TSPyV ST). The percent change in serum dependence doubling time (solid bars) was normalized (dashed bars) by glucose uptake compared to pLENTI control (H). MCPyV ST, HPyV7 ST, and TSPyV ST protein structures were determined and aligned using PyMol and black arrows denote the structurally dissimilar regions in protein structure (I). Each point represents mean ± the standard error of independent triplicates. One-way ANOVA * p < 0.05; ** < 0.01; *** p < 0.001, significantly different from pLENTI.

Consistent with TSPyV and HPyV7 not being found to be associated with cancer, expression of HPyV7 and TSPyV ST proteins in Rat-2 cells failed to significantly induce anchorage-independent growth above control cells (Fig. 2B and C). Furthermore, the metabolic rate, as measured by glucose uptake, of HPyV7 and TSPyV ST were lower than that of MCPyV ST (Fig. 2D). Interestingly, although the proliferation rate, doubling time, and serum dependence of HPyV7 ST were more similar to control cells, TSPyV ST induced a cellular proliferation rate, doubling time, and serum dependence similar to that of MCPyV ST, further demonstrating that transformation is a combination of many cellular abnormalities that are not necessarily connected (Fig. 2E, F, G, and H).

Together, our results indicate that MCPyV ST can uniquely induce anchorage-independent growth, increase glucose uptake and proliferation rate, and reduce doubling time and serum dependence when compared to HPyV7 ST; however, although TSPyV ST was incapable of inducing anchorage-independent growth and increased glucose uptake, it was able to increase cellular proliferation and reduce doubling time and serum dependence. In an effort to determine the domain(s) responsible for the unique transforming capabilities of MCPyV ST, an amino acid alignment of MCPyV, TSPyV, and HPyV7 ST was performed (Supplementary Figure 1). Unfortunately, although all evolutionarily related ST proteins, they contained exceptionally different amino acid sequences that made it extremely difficult to identify MCPyV ST unique domains possibly responsible for transformation. Despite dissimilar amino acid sequences, MCPyV, TSPyV, and HPyV7 ST proteins were found to have very similar protein structures upon protein structure alignment (Fig. 2I). Consistent with the common T region being hypothesized to not contain the domain(s) necessary for MCPyV ST transformation (Fig. 1I), the common-T structure of the transforming MCPyV ST, and non-transforming TSPyV and HPyV7 ST were very similar. However, despite their similarity, MCPyV, TSPyV, and HPyV7 ST contained structurally dissimilar loops in the ST-unique region, consistent with the previous hypothesis of the transforming domains being found in the ST-unique region (Fig. 1I). Together, these data suggest that the unique transforming capabilities of MCPyV ST may be a result of the activities of the structurally dissimilar loops in the unique region.

### MCPyV ST binds to many nuclear cellular proteins

Previous studies of the MCPyV viral life cycle and MCPyV ST-mediated cellular transformation have identified many MCPyV ST cellular interactors (13,15,16,34,36,40–45). Interestingly, when reviewing these findings, we found that 74% of the previously identified MCPyV ST cellular interactors are nuclear proteins (Fig. 3A). These findings were unexpected, as the amino acid sequence of MCPyV ST does not contain a known, canonical NLS, nor do many cellular localization programs such as PSORTII, seqNLS, and BaCelLO predict MCPyV ST to exhibit nuclear localization (Fig. 3B and C). Although MCPyV LT has been found to localize to the nucleus as a result of its NLS, the NLS of LT is found in the LT unique region, and mutation of the common-T region of LT did not affect its nuclear localization (Fig. 3D) (46). Therefore, in addition to transformation, the potential nuclear localization of MCPyV ST may also be due to the activity of a domain found in the ST unique region.

**Figure 3.**
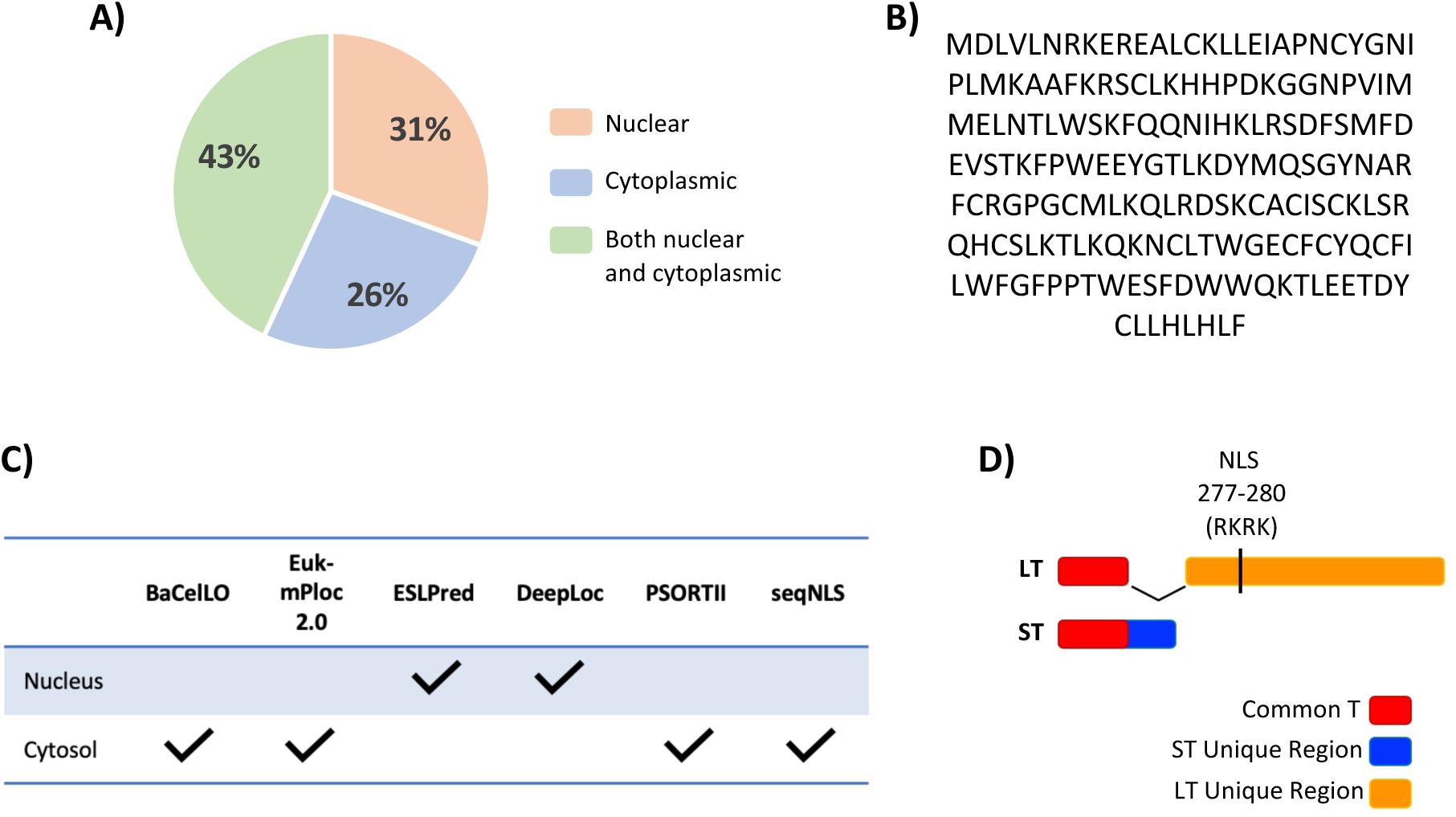
MCPyV ST binds to many nuclear cellular proteins. The localization of known MCPyV ST cellular interactors were recorded, with 31% being nuclear (orange), 26% being cytoplasmic (blue), and 43% that are both cytoplasmic and nuclear (A). The amino acid sequence of MCPyV ST contains no known canonical NLS (B). Online amino acid and protein subcellular localization predictor tools were utilized to predict the localization of MCPyV ST (C). The location of the MCPyV LT-t NLS is found within the MCPyV LT-t unique region (orange), but no known NLS is found within the common-T region (red) or ST unique region (blue) (D).

### MCPyV ST uniquely localizes to the nucleus independent of size

In order to determine whether MCPyV ST can localize to the nucleus in the absence of a canonical NLS, subcellular fractionations (SCF) were performed. Rat-2 cells transduced with pLENTI MCPyV ST were fractionated into cytoplasmic, nuclear, and membrane fractions, with MCPyV ST being found to localize to both the cytoplasmic and nuclear subcellular compartments (Fig. 4A). Although a small amount of MCPyV ST protein was detected in the membrane fraction, it is consistent with the level of HSP90 cytoplasmic contamination within the membrane fraction, and is likely a false signal. To confirm this finding in another cell line, 293As transfected with pMTBS also identified MCPyV ST to localize to the cytoplasmic and nuclear fractions, with membrane localization again likely a result of cytoplasmic contamination (Fig. 4B). Of note, it was consistently found that SCF experiments in 293A cells frequently led to more contamination than Rat-2 cells; however, the results from both cell lines confirmed both MCPyV ST cytoplasmic and nuclear localization.

**Figure 4.**
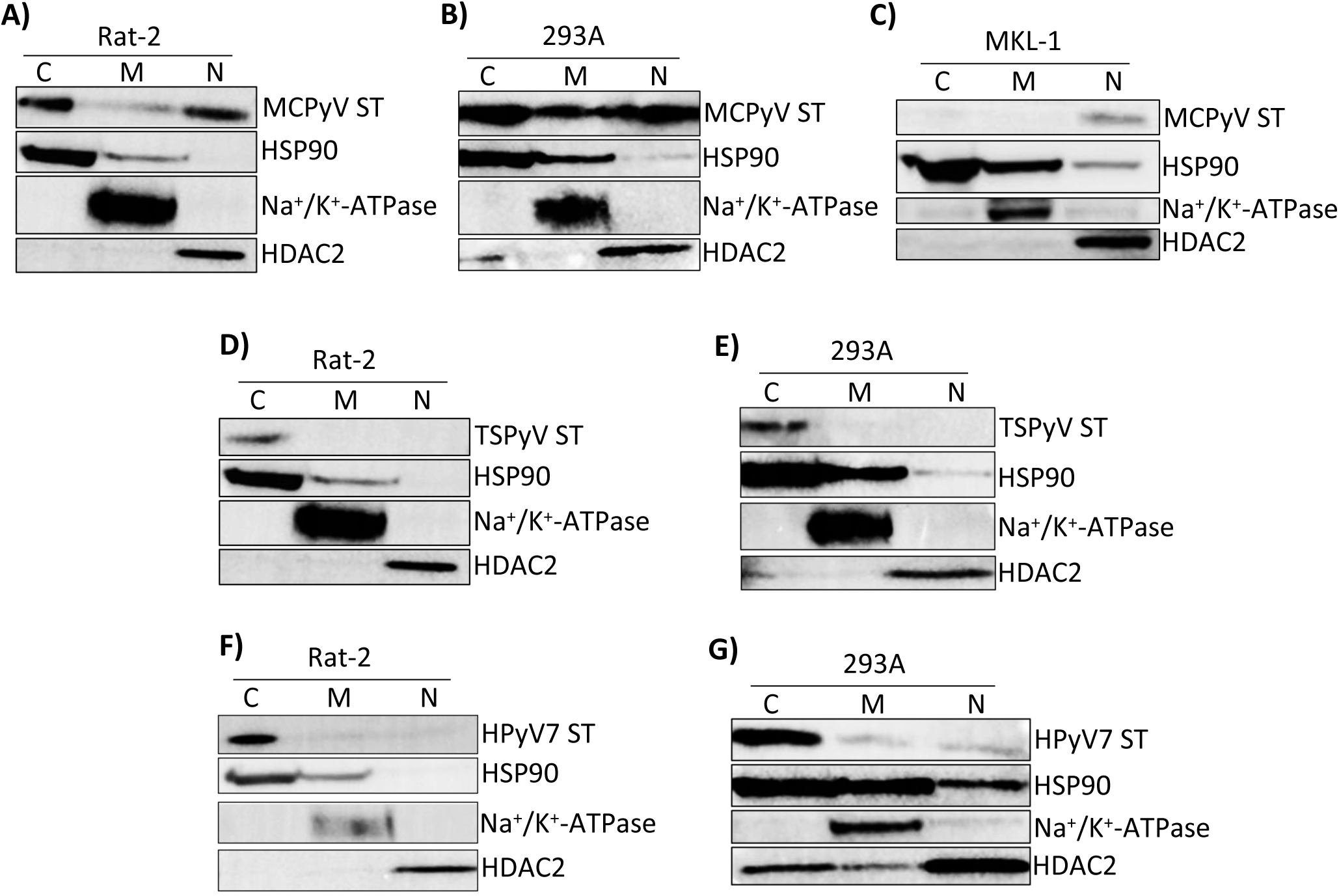
MCPyV ST uniquely localizes to the nucleus independent of size. Rat-2 (A) and 293A (B) cells were transduced and transfected, respectively, with MCPyV ST and a subcellular fractionation was performed to isolate proteins for the cytoplasmic (C), membrane (M), and nuclear (N) fractions. A subcellular fractionation for MCPyV ST was also performed on the VP-MMCC cells, MKL-1 (C). Rat-2 and 293A cells were also transduced and transfected, respectively, with TSPyV ST (D and E) and HPyV7 ST (F and G), followed by subcellular fractionation. HSP90 was used as a cytoplasmic control, NA^+^/K^+^-ATPase was used as a membrane control, and HDAC2 was used as a nuclear control.

To confirm these findings in a physiologically relevant cell line, an SCF was performed in MKL-1 cells, a virus-positive MCC cell line. Interestingly, in a VP-MCC cell line, endogenous MCPyV ST localized exclusively to the nucleus (Fig. 4C). This finding was consistent with exogenous MCPyV ST nuclear localization in Rat-2 and 293A cells, but inconsistent with shared cytoplasmic localization.

As shown previously, MCPyV ST is uniquely transforming when compared to other human, skin-tropic polyomaviruses such as TSPyV and HPyV7 ST (Fig. 2). To determine whether MCPyV ST is also unique in its ability to localize to the nucleus despite the absence of a canonical NLS, SCFs were performed on Rat-2 and 293A cells transfected with pCS2 TSPyV ST or pCS2 HPyV7 ST. Interestingly, both TSPyV ST (Fig. 4D and E) and HPyV7 ST (Fig. 4F and G) exclusively localized to the cytoplasm in both Rat-2 and 293A cells. These data suggest that MCPyV ST is unique in both its transforming capacity and nuclear localization. Furthermore, the domain(s) responsible for the unique transformation and nuclear localization of MCPyV ST may be found in the structurally dissimilar loops of the ST unique region (Fig. 2I).

### MCPyV ST nuclear localization is important for anchorage-independent growth, increased metabolism, and decreased serum dependence of MCPyV ST, but is expendable for proliferation rate and doubling time

In order to determine whether nuclear localization of MCPyV ST is necessary for cellular transformation, both a cytoplasmic and nuclear sequestered MCPyV ST were created by adding the NLS from SV40, or the nuclear export signal (NES) from Mitogen-Activated Protein Kinase Kinase (MAPKK), to MCPyV ST (NLS-ST and NES-ST, respectively). Importantly, localization domains were added to the N-terminus of MCPyV ST to ensure the furthest distance from the likely transforming domains within the structurally dissimilar loops of the MCPyV ST unique region in the C-terminus (Fig. 1I and 2I). Furthermore, localization domains were added with the addition of a flexible linker to decrease the likelihood of the added domain interfering with the structure and/or functions of MCPyV ST. To ensure the expression and stability of the MCPyV ST localization mutants, a confirmation western blot was performed and found the localization mutants and wild-type MCPyV (WT-ST) to all be expressed; however, protein densitometry revealed NLS-ST and NES-ST to be expressed at 41% and 59% of WT-ST, respectively. SCF experiments in Rat-2 cells confirmed the appropriate localization of WT-ST and the localization mutants (Fig. 5B). Similar to WT-ST, NLS-ST retained localization to both the nucleus and the cytoplasm, with a higher prevalence of nuclear localization. More importantly, NES-ST localized exclusively to the cytoplasm, necessary to assess the importance of nuclear localization in MCPyV ST-mediated cellular transformation.

**Figure 5.**
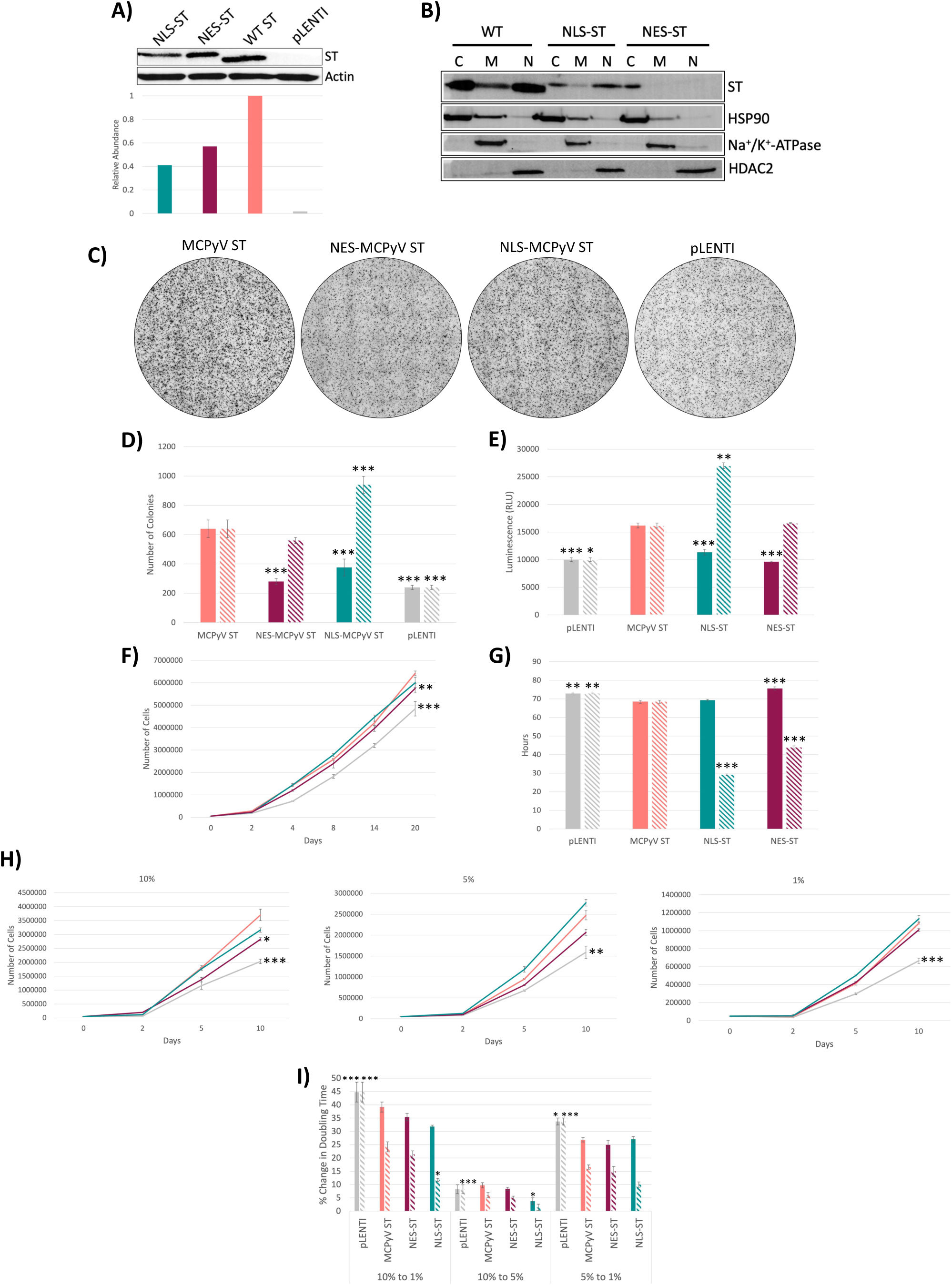
MCPyV ST nuclear localization is important for anchorage-independent growth, increased metabolism, and decreased serum dependence of MCPyV ST, but is expendable for proliferation rate and doubling time. Rat-2 cells were transduced with MCPyV ST (ST-WT), a cytoplasmic sequestered MCPyV ST (NES-ST), a nuclear sequestered MCPyV ST (NLS-ST), or pLENTI control, and protein expression was confirmed with protein densitometry (A). A subcellular fractionation was performed to verify localization of the wildtype ST and localization mutants. Subcellular fractionations included cytoplasmic (C), membrane (M), and nuclear (N) fractions, with HSP90 acting as a cytoplasmic control, NA^+^/K^+^-ATPase as a membrane control, and HDAC2 as a nuclear control (B). Various transformation assays were performed, including soft agar assays (C and D), glucose uptake (E), proliferation rate (F), doubling time (G), and serum dependence (H) (grey – pLENTI control, coral – MCPyV ST, maroon – NES-ST, teal – NLS-ST). The number of soft agar colonies (D), glucose uptake (E), and doubling time (G) were normalized based on protein levels determined by densitometry (solid bars – unnormalized, dashed bars – normalized) (A). The percent change in serum dependence doubling time (solid bars) was normalized (dashed bars) by densitometry determined protein levels and glucose uptake compared to wildtype MCPyV ST (I). Each point represents mean ± the standard error of independent triplicates. One-way ANOVA * p < 0.05; ** < 0.01; *** p < 0.001, significantly different from MCPyV ST.

To assess the effect of localization on MCPyV ST-mediated cellular transformation, various transformation assays were performed comparing WT-ST, NLS-ST, and NES-ST, followed by normalization based on protein densitometry. NLS-ST was capable of forming colonies above WT-ST post-normalization, whereas NES-ST also led to colony formation, but in fewer numbers (Fig. 5 C and D). Normalized glucose uptake of NLS-ST was significantly higher than that of WT-ST, whereas the glucose uptake of NES-ST was similar to that of WT-ST, suggesting nuclear localization of MCPyV ST may influence the ability of MCPyV ST to increase cellular metabolism (Fig. 5E). The proliferation curves of WT-ST and both localization mutants were significantly increased when compared to the empty pLENTI transduced controls, but were not significantly different from each other (Fig. 5F). However, the protein-level normalized doubling times of NLS-ST and NES-ST were both significantly faster than WT-ST, with NLS-ST having the fastest doubling time (Fig. 5G).

No significant differences were observed when comparing the proliferation rate curves of WT-ST compared to the localization mutants at differing serum concentrations, likely as a result of the differing protein levels and metabolic rates of each ST (Fig. 5H). Therefore, the percent change in doubling time between differing serum concentrations was calculated and normalized by both the protein levels (Fig. 5A) and glucose uptake rates (Fig. 5E) of each ST (Fig. 5I). Together, WT-ST and NES-ST were each found to have similar changes in the percent doubling time between 10% and 1% serum containing media, whereas NLS-ST was found to have a significantly lower percent change in the doubling time between 10% and 1% serum-containing media. Together, these data suggest nuclear localization of ST is largely responsible for the ability of MCPyV ST to proliferate in the absence of serum despite having an increased metabolic rate, as inferred by an increased glucose uptake. In conclusion, these data suggest that cellular proliferation and doubling time may be a combined result of both nuclear and cytoplasmic localization, whereas increased anchorage independent growth, metabolism, and decreased serum dependence may be influenced by the nuclear localization of MCPyV ST.

## Discussion

The mechanism(s) of MCPyV-mediated cellular transformation and tumorigenesis have been investigated since the discovery of MCPyV in 2008; however, many of these early analyses were hindered by using the approach of comparing the similarities between the human oncogenic polyomavirus MCPyV, and the simian oncogenic virus SV40. The findings of the current study and of others that ST is the dominant transforming protein of MCPyV is opposite to SV40, in which ST plays only a supportive or accessory role, with LT being the dominant transforming protein (47). Similarity comparisons between the dissimilar SV40 and MCPyV are responsible for many years spent investigating MCPyV ST binding to PP2A and Fbw7, which have been found to be irrelevant for transformation by MCPyV ST (40,41). Furthermore, we found MCPyV ST to be uniquely transforming among skin-tropic human polyomaviruses ST antigens, consistent with MCPyV being the only human polyomavirus clearly associated with cancer. Together, these data refute the method of identifying mechanisms of MCPyV transformation through similarity comparisons to SV40, and support the alternative dissimilarity approach of investigating the differences between MCPyV ST and the ST of other non-oncogenic human polyomaviruses to identify the unique mechanisms of MCPyV ST mediated transformation.

The dissimilarity approach can also be applied when comparing MCPyV ST and LT-t, as they share a common T region, have their own unique regions, and accomplish dissimilar patterns of transformation. Through this approach, we were able to narrow down the location of the domain(s) responsible for MCPyV ST transformation to the ST unique region. Although transformation assays performed on Rat-2 cells expressing the ST unique region alone may be able to confirm this finding, it is likely that the absence of common-T may influence the structure of the ST unique region, and it is possible that domain(s) within the common-T region may be necessary to facilitate the activities of the ST unique region.

Although the discovery that MCPyV is uniquely oncogenic among human polyomaviruses was discouraging at the genesis of MCPyV oncogenesis research, the dissimilarity approach exploited it for our benefit. Comprehensive transformation assay analysis of TSPyV ST and HPyV7 ST identified that although the ST of HPyV7 and TSPyV were not as robustly transforming as MCPyV ST, TSPyV ST was capable of achieving some properties of cellular transformation, further supporting the fact that cellular transformation is a collection of properties rather than a single entity, and that the ability to induce several transforming properties may translate to being able to accomplish tumorigenesis *in vivo.* Furthermore, structural alignments allowed us to further narrow down the unique transforming domains of MCPyV ST to the structurally dissimilar loops consistent with the unique region of MCPyV ST. It is currently hypothesized that the similar transforming capabilities of MCPyV and TSPyV ST may reside in domains shared by MCPyV and TSPyV ST, but not found in HPyV7 ST. Alternatively, TSPyV ST and MCPyV ST may accomplish similar methods of cellular transformation through distinct mechanisms.

The identification of MCPyV ST binding to several nuclear proteins suggested nuclear localization despite the absence of a canonical NLS in MCPyV ST. Surprisingly, subcellular fractionation confirmed nuclear localization of MCPyV ST in various cell lines, whereas the non-oncogenic TSPyV and HPyV7 ST proteins were confined to the cytoplasm, further proving the usefulness of the dissimilarity hypothesis. Although MCPyV ST is under the 30kDa size necessary for passive diffusion through the nuclear pore, the finding that 30kDa TSPyV and HPyV7 ST proteins are cytoplasmic confirms that MCPyV ST localization to the nucleus is regulated (48). Interestingly, MCPyV ST exhibited both nuclear and cytoplasmic localization in transduced Rat-2 and 293A cells, yet it was exclusively nuclear in VP-MCC cells. Although it is possible that the differences in MCPyV ST localization between the cell lines may be an artifact of differing cell lines, hypotheses to explain these differences include the following: First, it is possible that the mechanism of MCPyV ST nuclear translocation is saturated in Rat-2 and 293A cells, overexpressing exogenous ST, leading to spillover into the cytoplasm. Second, the presence of MCPyV LT-t in MKL-1 cells, but not Rat-2 and 293A cells, may enhance the efficiency of MCPyV ST nuclear localization. Although both hypotheses require further experimentation, it is clear from these experiments that MCPyV ST is capable of uniquely localizing to the nucleus despite the absence of a canonical NLS.

It can be hypothesized that the noncanonical NLS or domain responsible for MCPyV ST nuclear localization is found in the MCPyV ST unique region, as previous investigation into MCPyV LT-t nuclear localization identified the common T region to not be responsible for nuclear localization (46). As the function of an NLS is dependent on both structure and function, it is possible that the MCPyV ST NLS is located in the highly dissimilar amino acid sequence or the structurally dissimilar loops between MCPyV, TSPyV, and HPyV7 ST.

Although most of the attention regarding MCPyV-mediated oncogenesis has been directed towards ST, it is clear that the large T antigen is also important due to its necessity for the viability of MCC cells and its ability to influence cellular proliferation and doubling time, despite not inducing anchorage-independent growth, metabolism, and serum dependence. Due to the sharing of the common-T region between LT and ST, the domain(s) responsible for inducing proliferation rate and doubling time may be found in the common T region; however, it is more likely that ST and LT-t accomplish this through different mechanisms attributed to their common-T region, as they affected Rat-2 cells dissimilarly, despite similar protein expression levels. It is likely that perturbation of Rb by the Rb binding domain of MCPyV LT-t contributes to these phenotypes; however, this has not yet been directly assessed.

Finally, the export of MCPyV ST to the cytoplasm did not ablate all properties of cellular transformation, suggesting a potential role of MCPyV in both the cytoplasm and nucleus. However, it is not currently understood how this finding translates to the observation of MCPyV ST being exclusively nuclear in VP-MCC cells, and therefore warrants further investigation. Furthermore, although normalized NES-ST was still capable of anchorage-independent growth, increased glucose uptake, and reduced serum dependence, NLS-ST normalization led to anchorage-independent growth, glucose uptake, and serum dependence significantly higher than WT-ST and NES-ST. Although protein levels of WT-ST and NLS-ST differed, the ratio of WT-ST and NLS-ST in the cytoplasmic and nuclear compartments were similar. Therefore, it is of great interest why NLS-ST was capable of increasing transformation above WT-ST in some assays but not others.

## Conclusions

Together, it has been found that elucidating mechanisms of transformation guided by a dissimilarity approach may prove useful in identifying otherwise hidden, unique mechanisms of MCPyV oncogenesis, such as the unique nuclear localization of MCPyV ST. Cellular transformation was found to be multi-variable and not simply a positive or negative characteristic of viral proteins, consistent with our understanding of the role of viral proteins in viral replication. Further investigation into the mechanisms and role of MCPyV nuclear localization in transformed and MCC cells will further enhance our understanding of the natural viral life cycle and MCPyV-mediated oncogenesis necessary for basic virology and the treatment of MCC.

## List of Abbreviations

MCC: Merkel Cell Carcinoma
MCPyV: Merkel Cell Polyomavirus
HPyV7: Human Polyomavirus 7
TSPyV: Trichodysplasia Spinulosa Polyomavirus
HPyV: Human Polyomavirus
NES: Nuclear Export Signal
NLS: Nuclear Localization Signal
Tag: Tumor Antigens
ST: Small Tumor Antigen
LT-t: Truncated Large Tumor Antigen
ALTO: Alternate Large T Open Reading Frame
ER: Early Region
LR: Late Region
VP-MCC: Virus Positive Merkel Cell Carcinoma
FBS: Fetal Bovine Serum
SCF: Subcellular Fractionation
SAA: Soft Agar Assay
SV40: Simian Virus 40

## Declarations

## Ethics approval and consent to participate

Not applicable.

## Consent for publication

Not applicable.

## Availability of data and materials

The datasets supporting the conclusions of this article are included within the article and supplementary materials.

## Competing interests

The authors declare no conflict of interest.

## Funding

Finanpcial support was provided by the Stetson University Health Sciences Department and Cell Biology Education Consortium.

## Author Contributions

K.T., I.T., and C.D. facilitated conceptualization, investigation, data curation, writing of the original draft, review and editing, and visualization. M.S. performed formal analysis and review and editing. K.D. performed conceptualization, methodology, validation, investigation, writing of original draft and review and editing, visualization, supervision, and funding acquisition.

## Acknowledgments

The plasmids and lentiviruses used in these studies, and the modeling of ST structures were made in the laboratory of Dr. Denise A. Galloway at the Fred Hutchinson Cancer Center when Dr. Dye was a graduate student. This work was supported by R35-CA209979 to DAG and by T32-AIO83203 to KND. Furthermore, we would also like to thank Richard Wang (University of Texas) for sharing Rat-2 cells, and Dr. Patrick Moore and Dr. Yuan Chang (University of Pittsburgh) for sharing MKL-1 cells, and Dr. James DeCaprio (Harvard) for donating Ab5.

---

**Supplementary Figure 1:**
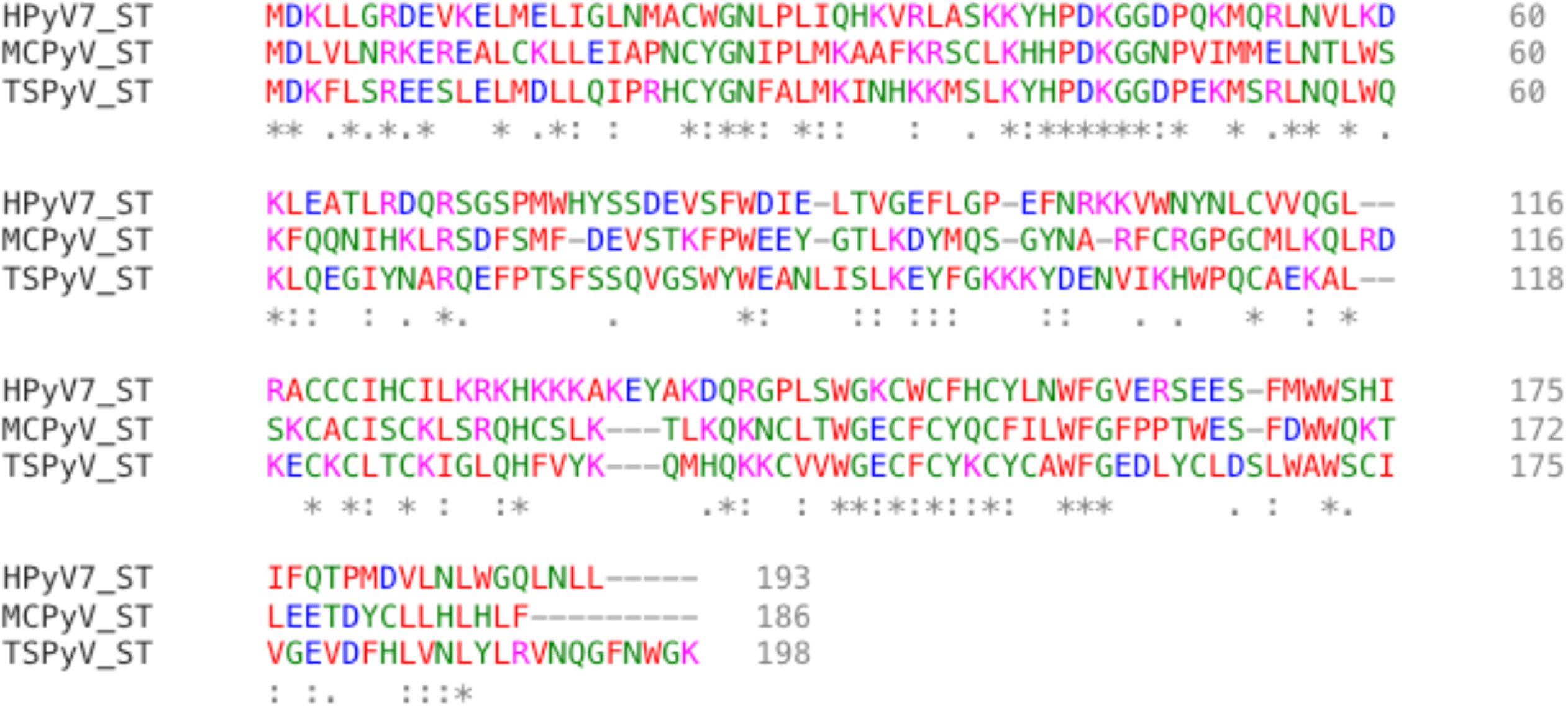
An amino acid alignment of MCPyV ST, HPyV7 ST, and TSPyV ST created using ClustalOmega.com.

## Notes

### Competing Interest Statement

The authors have declared no competing interest.

### Summary of Updates

Changes were made to figure 3.

